# Estimating the reproduction number, *R*_0_, from agent-based models of tree disease spread

**DOI:** 10.1101/2023.08.03.551748

**Authors:** Laura E Wadkin, John Holden, Rammile Ettelaie, Melvin J Holmes, James Smith, Andrew Golightly, Nick G Parker, Andrew W Baggaley

## Abstract

Tree populations worldwide are facing an unprecedented threat from a variety of tree diseases and invasive pests. Their spread, exacerbated by increasing globalisation and climate change, has an enormous environmental, economic and social impact. Computational agent-based models are a popular tool for describing and forecasting the spread of tree diseases due to their flexibility and ability to reveal collective behaviours. In this paper we present a versatile agent-based model with a Gaussian infectivity kernel to describe the spread of a generic tree disease through a synthetic treescape. We then explore several methods of calculating the basic reproduction number *R*_0_, a characteristic measurement of disease infectivity, defining the expected number of new infections resulting from one newly infected individual throughout their infectious period. It is a useful comparative summary parameter of a disease and can be used to explore the threshold dynamics of epidemics through mathematical models. We demonstrate several methods of estimating *R*_0_ through the agent-based model, including contact tracing, inferring the Kermack-McKendrick SIR model parameters using the linear noise approximation, and an analytical approximation. As an illustrative example, we then use the model and each of the methods to calculate estimates of *R*_0_ for the ash dieback epidemic in the UK.

## 1. Introduction

The loss of biodiversity due to the spread of tree diseases and invasive pests within forest ecosystems has an enormous environmental, economic, and social impact worldwide [1, 2]. This threat has been escalating rapidly due to increasing globalisation resulting in a greater number of accidental imports and climate change creating a more favourable environment for many pests and pathogens.

Mathematical and computational models are powerful tools for deepening our understanding of the fundamental behaviours of different pests and pathogens, as well as describing and forecasting their future spread [3, 4, 5, 6]. Agent-based models, where each ‘agent’ represents an individual tree (or group of trees) with properties governed by a set of probabilistic rules, have the advantage of providing information about the whole system (i.e., the macroscale and collective behaviour) from modelling the individual (microscale) behaviour and are ideal for exploring spatial patterning. Previous compartmental agent-based models of tree disease have considered a square lattice representing susceptible trees (or areas of trees) with pathogen spread stochastically through nearest neighbour interactions [7]. Although suitable for some fungal diseases [8, 7], many tree diseases are spread through airborne pathogens on longer scales and are not suitable for nearest neighbour contact spread [9, 10]. Thus, here we adopt an agent-based model with a dispersal kernel representing the spatial dispersal of the pathogen or pest which results in the probability of infection.

The infectivity of a disease can be quantified by the basic reproduction number, *R*_0_, defined as the total number of expected secondary infections arising from one newly infected individual introduced into a fully susceptible population. It helpfully defines the threshold between epidemic (*R*_0_ > 1), and containment (*R*_0_ < 1), and can be used as a comparative parameter for diseases with differing behaviours and spread mechanisms. In many mathematical models (including the Kermack-McKendrick SIR model considered in this text) *R*_0_ can be used to determine the steady-state conditions of the disease or pathogen density in a host population [11, 12, 13, 14, 15]. It is possible to calculate *R*_0_ from knowledge of the pathogen’s life-cycle and its interactions with the host plant [15], however this information is not always readily available. Previous work has considered estimating *R*_0_ through a spatially-explicit population dynamic model for wheat stripe rust [16], the SIR model with a time-varying infestation rate for the oak processionary moth pest [17] and through stochastic epidemiological landscape models for sudden oak death [18].

In this paper, we present a versatile agent-based model with a Gaussian infectivity kernel (referred to henceforth as the ABM) and explore several methods for estimating the basic reproduction number from this model, including contact tracing, parameter inference for a SIR model and an analytic approximation. We then take an illustrative case-study of the UK ash dieback epidemic to compare these methods. We outline the ABM, the compartmental SIR model with accompanying parameter inference methodology, and the analytic approximation of *R*_0_ in Section 2, present the results in Section 3, and discuss the findings in Section 4.

## 2. Methods

In this section we introduce the basic reproduction number *R*_0_ (Section 2.1), present the ABM for a generic tree disease (Section 2.2), summarise the stochastic SIR model (Section 2.3), outline the statistical methodology for parameter inference of the stochastic SIR model (Section 2.4), and derive an analytical approximation of *R*_0_ for the ABM (Section 2.5).

### 2.1. The reproduction number R_0_

The reproduction number is a key parameter quantifying the spread of a disease. The basic reproduction number, *R*_0_, describes the expected number of secondary infections produced by one primary infected tree, over their infectious period. Some of the methods presented in Section 3 estimate the basic reproduction number at every time-step of the simulation, which we will refer to as the instantaneous basic reproduction number, *R*_0_(*t*). We will also consider the instantaneous effective reproduction number, calculated at a time *t* as *R*(*t*) = *R*_*0*_*S*(*t*)*/S*(0), which takes into account that as a disease progresses the number of susceptibles, *S*(*t*), is decreasing and thus limiting disease transmission.

### 2.2. The agent-based model

We consider individual trees as points, randomly distributed at a density *ρ* within a bounded square of *L× L*. All trees are classified as being in one of three states: susceptible (*S*), infected (*I*) or removed (*R*^*†*^) (a compartmental SIR model as described in Section 2.3 [11]). Susceptible trees have yet to be infected and are at risk, infected trees currently have the disease and are infective to surrounding susceptible trees during their infectious period *T*_I_, and removed trees have previously been infected, but are no longer infectious. Trees transition through the states *S* → *I* → *R*^*†*^ with opportunities to transition in every iteration of an arbitrary discrete time-step which can be rescaled to handle different time courses.

We assume that susceptible trees at shorter distances from infected trees are more likely to become infected than those further away. Thus, infection spreads through the trees based on a spatial kernel, allowing a dependence on the distance between a susceptible and an infectious tree. The choice of this kernel is flexible, linking to the spatial dispersal of a pathogen and the probability of infection upon pathogen presence. A Gaussian kernel is commonly used, however, in some ecological cases other kernels may be more appropriate [19, 10]. For simplicity, here we apply a Gaussian kernel to capture the decay of infection probability with increasing distance between trees, but this is easily transferable to any other kernel function. In this case, the probability of a susceptible tree *S*_*i*_ becoming infected due to an infectious tree *I*_*j*_, separated by a distance *r*, in an arbitrary time-step of the ABM Δ*t*, is described by the function

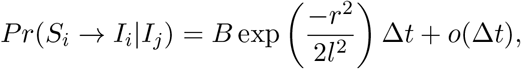

where *B* is an infectivity parameter, *l* is the length scale of the pathogen dispersal and *o*(Δ*t*)*/*Δ *t* → 0 as Δ*t* → 0. To make epidemics comparable as the transmission length scale varies, we scale the infectivity parameter, leading to the kernel

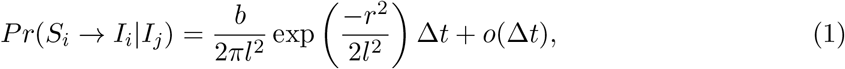

where Δ*t* = 1 and therefore *b/*(2*πl*^2^) = *B* takes a value between 0 and 1. Two example kernels of the above form with contrasting length scales are shown in Figure 1(a).

**Figure 1:**
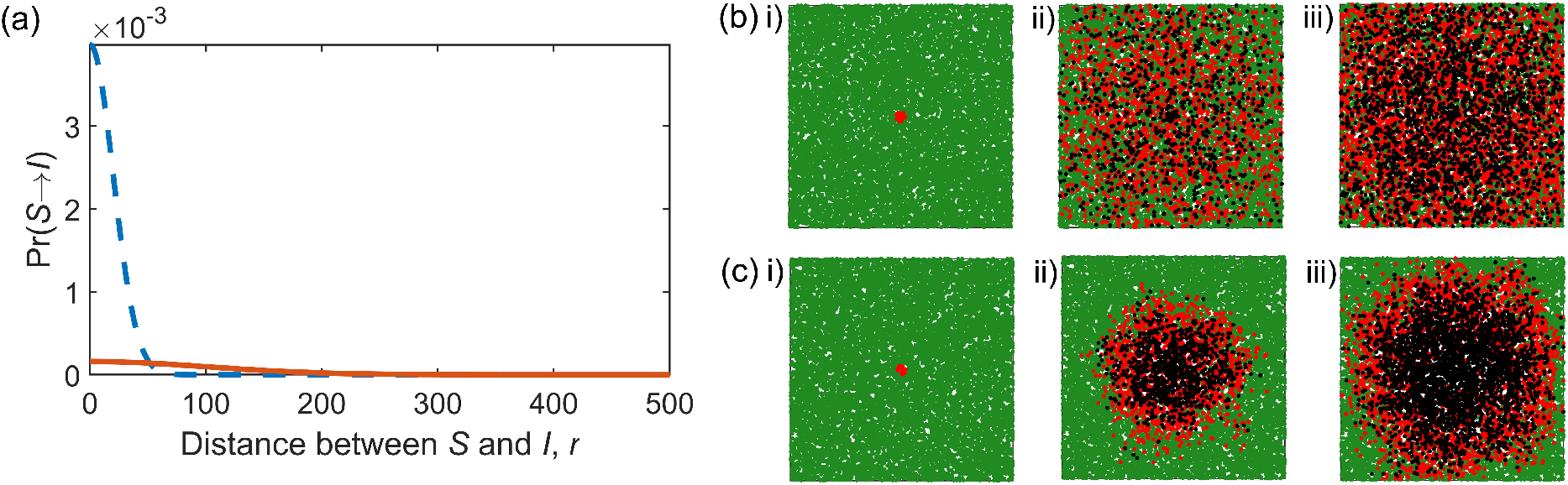
(a) Two example Gaussian infectivity kernels from (1), with *b* = 10, *l* = 100 (orange, long-range spread) and *b* = 10, *l* = 10 (blue dashed, short-range spread). The resulting infection dynamics are shown in (b) and (c) for the long and short-range spreads respectively, showing snapshots at i) *t* = 0, ii) 20% tree mortality and iii) 50% tree mortality. Both cases begin with *I*_0_ = 20 infected trees with an infectious period of *T*_*I*_ = 10 time-steps, amongst a population of susceptible trees with density *ρ* = 0.05 in a bounded box of size *L* = 500.

We also assume that a susceptible tree is more likely to become infected if it is surrounded by a greater number of infectious trees, thus we require the probability that tree *S*_*i*_ becomes infected through any of the infectious trees *I*_*j*_ (with *j* = 1, …, *N*_*I*_) and their respective probability of infecting *S*_*i*_, i.e., *p*_*ij*_. To avoid a lengthy union calculation, we calculate the probability of *S*_*i*_ remaining uninfected, through Π (1 – *p*_*ij*_). The infectious period is set by a fixed parameter *T*_*I*_. Once a tree has been infectious for *T*_*I*_ time-steps, it transitions into the removed (*R*^*†*^) category.

The simulation begins with an initial number of infected trees, *I*_0_, chosen stochastically from trees closest to the centre of the domain. In one time-step all the possible infections are assessed and the corresponding compartmental transitions are applied. Example snapshots from the model are shown for the two illustrative infectivity kernels with contrasting length scale parameters in Figure 1(b) and (c).

#### 2.2.1. Contact tracing through the ABM

The most direct way to calculate *R*_0_ in a computational setting is through the contact tracing of secondary infections throughout the simulation. This is straightforward if the disease dynamics are one-to-one, with a tree becoming infected due to a ‘contact’ with a single infectious tree. However, in the ABM described above, we assume infections can occur due to pressure from multiple sources through the infectious kernels surrounding each infected tree, as shown in Figure 2.

**Figure 2:**
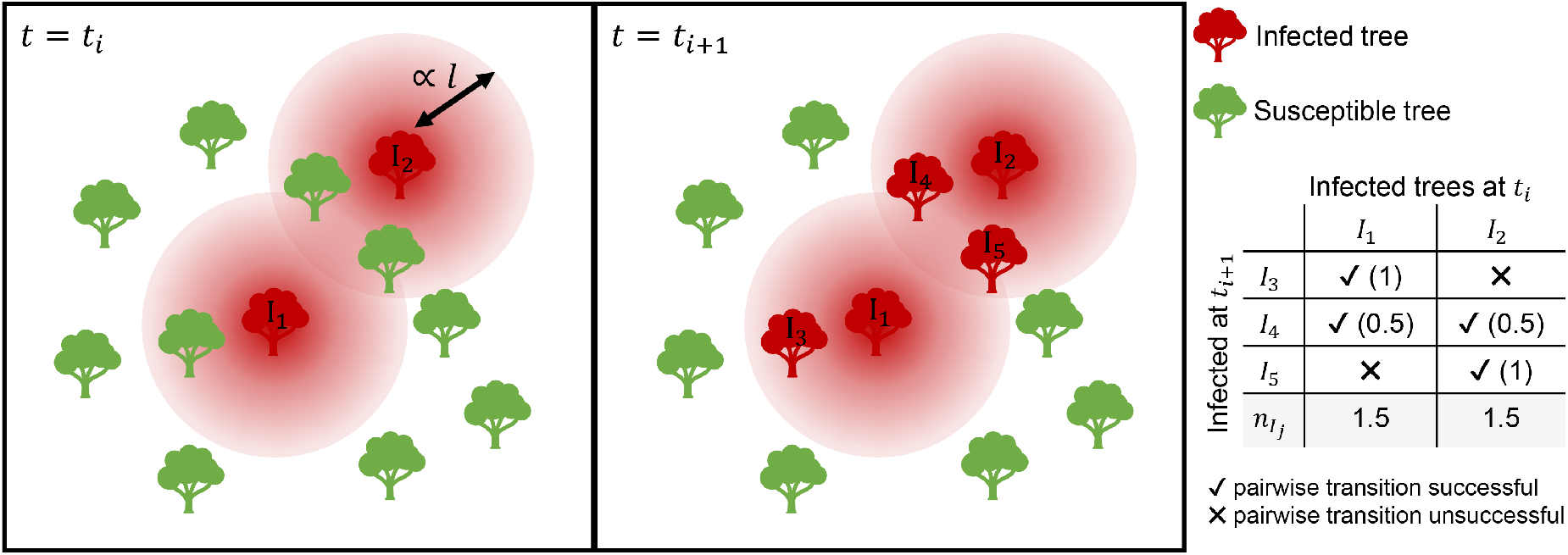
Schematic illustrating the number of secondary infections, *n*_*Ij*_, caused by two infected trees, *I*_1_ and *I*_2_. The circles around each of *I*_1_ and *I*_2_ at time-step *t* = *t*_*i*_ illustrate the surrounding infectivity kernel, with a length scale described by parameter *l*, and infectivity by *b*. For each tree within the infectious zone, the probability of transition is assessed as described in Section 2.2. For newly infected trees (i.e., *I*_3_, *I*_4_ and *I*_5_), the contributing infection pressures from previous infected trees (i.e., *I*_1_ and *I*_2_) are assessed independently, with proportional responsibility allocated to each previous infectious tree, as shown in the table.

In this case, after assessing the transition probabilities for all pairwise combinations of susceptible and infected trees, we can assign a resulting number of newly caused infections to each infectious tree that is proportional to the number of trees contributing towards the transition, as illustrated in Figure 2. For example, if tree *S*_1_ is successfully infected with transition probabilities positively assessed from tree *I*_1_ and *I*_2_, then both tree *I*_1_ and *I*_2_ are deemed to have caused 0.5 secondary infections for the transition of *S*_1_ into the infected category. These secondary infections can then be summed for each *I*_*i*_ after all *S*_*j*_ have been considered. This method conserves the total number of secondary infections caused, whilst allowing an insight into the individual tree contributions.

We can then estimate *R*(*t*) and *R*_0_ through this proxy-contact tracing method. The number of new secondary infections caused by each infected tree in each time-step can be averaged, giving the mean number of new infections in a time-step, 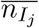. The instantaneous effective reproduction number is then 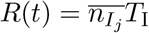 where *T*_I_ is the infectious period. For example, if in a single time-step there were 20 initially infected trees leading to 10 new infections, with a mean infection duration of 10 timesteps then 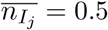 and *R*(*t* = 1) = 0.5 *×* 10 = 5, corresponding to the fact that at this stage of the dynamics there are five new infections (on average) from one initial infection over its infectious period. We present examples of this method as an estimate for *R*_0_ in Section 3.1.

### 2.3. The compartmental SIR model

We consider estimating the parameters for a standard SIR model to describe the ABM output as a method of calculating *R*_0_ (see Section 2.4 for inference details). The established formulation of a compartmental SIR model [20, 11] describes the rates of change of the populations in each compartment by

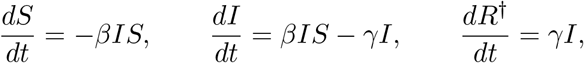

where *β* describes the rate of infection at which contact of one infected with one susceptible will result in infection due to pathogen dispersal (sometimes referred to as the effective contact rate), and *γ* describes the rate of removal due to a limited infectious period.

Since the ABM described above is inherently probabilistic, we will consider the more flexible stochastic SIR system (as opposed to the deterministic system) described by the Itô stochastic differential equation (SDE)

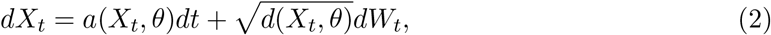

where *X*_*t*_ = (*S*_*t*_, *I*_*t*_)′ is the state of the system at time *t, θ* = (*β, γ*)′ is a vector of parameter values, and *dW*_*t*_ = (*W*_1,*t*_, *W*_2,*t*_)′ denotes a vector of uncorrelated standard Brownian motion processes. The SDE drift function *a*(*X*_*t*_, *θ*) and diffusion coefficient *d*(*X*_*t*_, *θ*) are given by

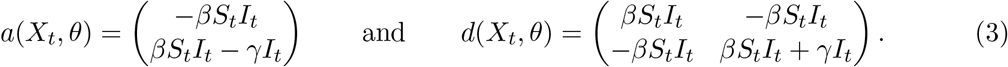

A derivation of the above can be found in [21]. The drift and diffusion functions are the infinitesimal mean and variance that match the most natural Markov jump process representation of the SIR model [22]. The conditions under which this leads to a reasonable approximation are also discussed in [22]. In this model, the basic reproduction number is given by *R*_0_ = *βN/γ*. In the next section we outline an inference scheme for estimating plausible values of *β* and *γ* (and thus *R*_0_) from the time-series output of the ABM.

Note that other variations of this model exist, allowing flexibility to capture the dynamics of different diseases, such as the SEIRS model [23] which includes additional exposed category, E, and feeds removed individuals back into the susceptible (S) category after a period of immunity. Similarly, it would be straightforward to adapt the corresponding categories in the ABM.

### 2.4. Bayesian inference

To perform Bayesian inference using the stochastic SIR model, (2) and (3), we apply the linear noise approximation (LNA) to obtain a tractable likelihood function, assume a Normal prior, and use a Markov chain Monte Carlo algorithm (see e.g., [24]) to sample values from the posterior distribution of the parameter *β*. We fix *γ* based on the set removal rate in the ABM, i.e., *γ* = 1*/T*_*I*_. Further details of this process are given in Appendix A.

### 2.5. An analytic approximation of R_0_

We consider an idealised, spatially explicit analytic expression of *R*_0_ for a single infectious individual. A full derivation is given in Appendix B. Firstly, consider a single infectious individual with infectious period *T*_*I*_, surrounded by susceptibles at varying distance *r* at a density *ρ*. We can estimate the number of new infections caused by the infected individual to be equal to the number of susceptibles at distance *r, S*(*t*) = 2*πrρ*, multiplied by the probability of infection at *r*, in this case described by (1). The expected number of new secondary infections from the initial infected individual at time step *t* (for 1 < *t* < *T*_*I*_) for all possible values of *r* is therefore

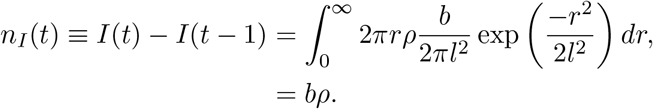

The cumulative sum of new secondary infections from the single infectious individual during a set time period 0 ≤ *t* ≤ *T*, denoted *N*_*I*_(*T*), could then be approximated as

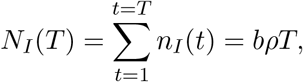

leading to a first approximation of *R*_0_ as the sum of secondary infections at the end of the infectious period *T*_*I*_:

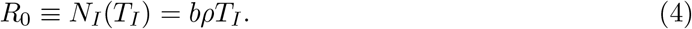

We refer to (4) as *R*_0_ approximation 1. However, this assumes that the density of the susceptibles remains fixed at each time step as the infection process progresses.

If we consider a large but finite domain of size *L*, we can introduce a time variant density taking into account the decreasing number of susceptibles due to the infection process, ρ(*t*), leading to *n*_*I*_(*t*) = *bρ*(*t*) with

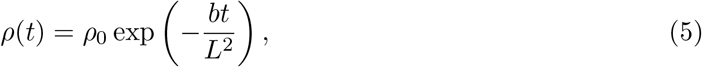

where *ρ*_0_ is the density at time *t* = 0. The cumulative sum of new secondary infections, during a set time period 0 ≤ *t* ≤ *T*, is then given by

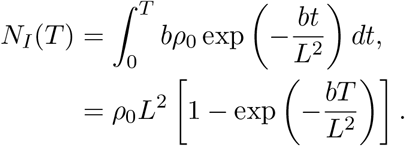

This is equivalent to noting that *N*_*I*_(*T*) is the overall change in the density of susceptibles, multiplied by the area, i.e., *N*_*I*_(*T*) = *L*^2^[*ρ*(*t* = 0) *− ρ*(*t* = *T*)]. The value of *R*_0_ is the sum of the new secondary infections across the whole infectious period,

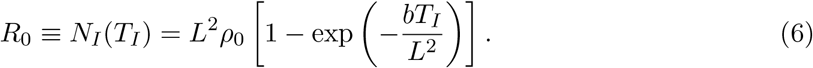

This can be considered as a second approximation to *R*_0_ where some reduction in the number of susceptibles has been taken into account. We refer to (6) as approximation 2. Note that this is an idealised scenario in which density reductions due to secondary and tertiary infections are neglected.

We can go a step further by considering the spatial influence of decreasing susceptibles. Due to the spatial infection kernel, we expect the density to vary both spatially (corresponding to the choice of infection kernel governing the spatial properties of the infection process) and temporally. We therefore consider a variant density of the form

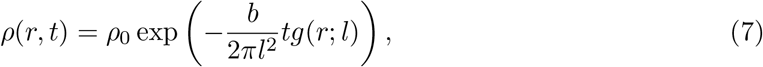

where *ρ*_0_ is the density at *t* = 0 and *g*(*r*; *l*) is a kernel equivalent to the infection kernel in (1). This density function, (7), is shown in Figure 3 for infection parameters *b* = 10 and *l* = 10 to illustrate its impact at short infection length scales. As with density (5) above, this function only considers the density reduction due to the secondary infections caused by the initial infected individual. The same approach could be taken for a different choice of infection kernel.

**Figure 3:**
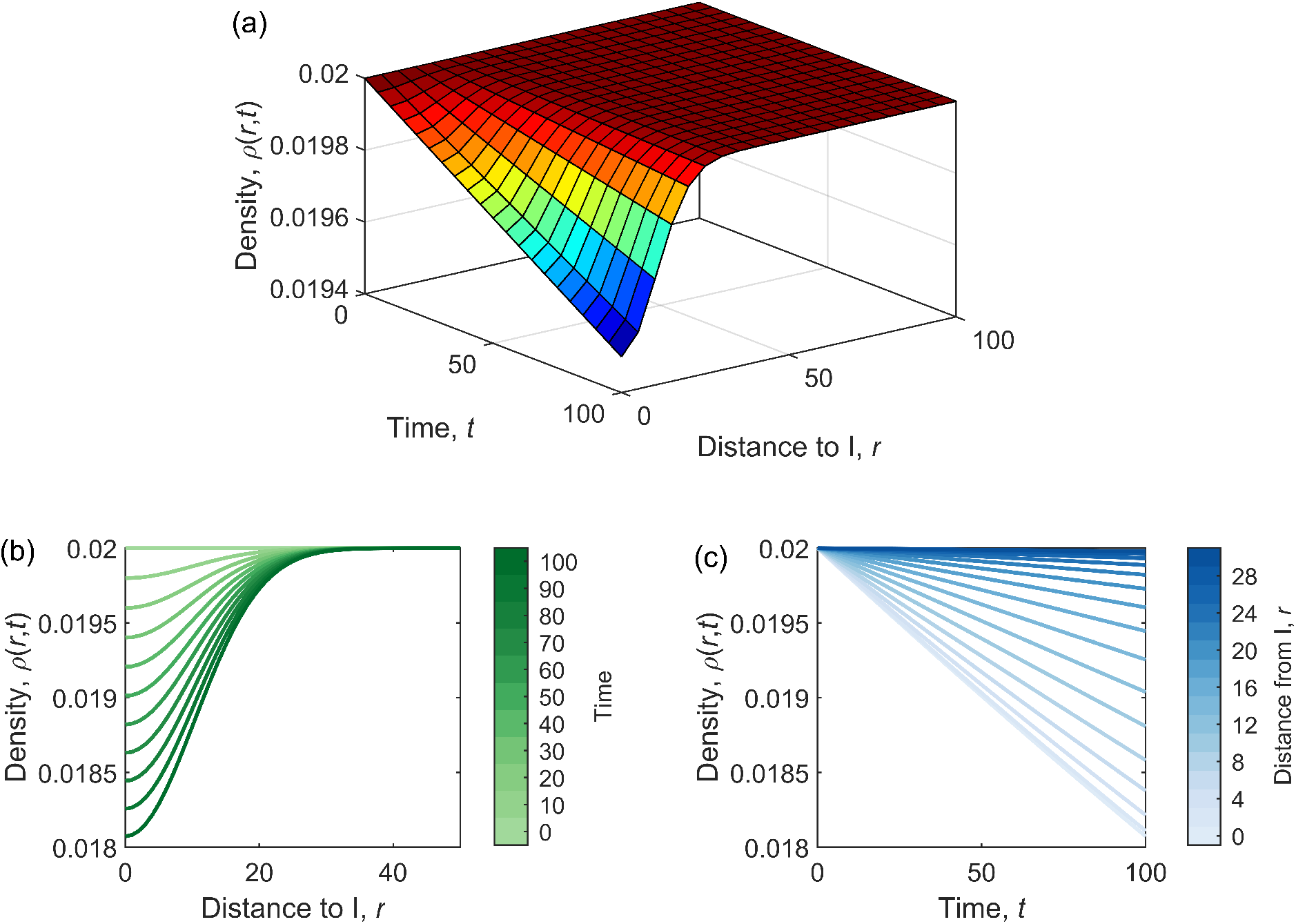
The analytic density function *ρ*(*r, t*) given by (7) for the idealised scenario of one infectious individual surrounded by susceptibles at an initial density of *ρ*_0_ = 0.02 and infection parameters *b* = 10 and *l* = 10 showing (a) the 3D surface *ρ*(*r, t*), (b) the changing density with distance from the initial infected and (c) the changing density with time.

As above, the total number of new secondary infections from the primary infection during a set time period, 0 ≤ *t* ≤ *T*, at a certain distance *r, N*_*I*_(*r, T*), can be calculated from the overall change in density

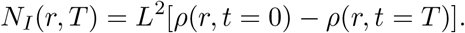

For all possible values of *r* this becomes

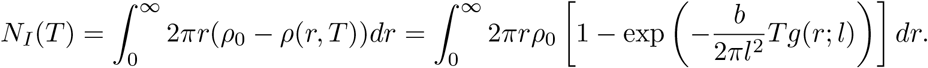

Although the above is difficult to solve directly, it can be integrated by performing a series expansion on the exponential term and integrating on a term-by-term basis, leading to

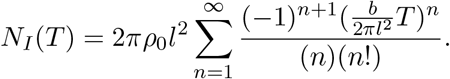

If *b/*2*πl*^2^ and *T* are small, the first order term is sufficient to approximate *R*_0_ as a linear function of *T*. This is confirmed by the numerical simulations in Section 3.3. Evaluating the summation gives

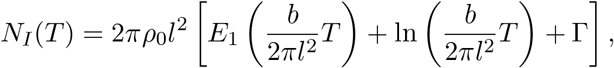

where the function *E*_1_(*x*) is the mathematically well studied exponential function 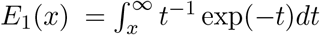 and Γ is the Euler-Mascheroni constant *≈* 0.57721. The estimated value of the basic reproduction number is therefore

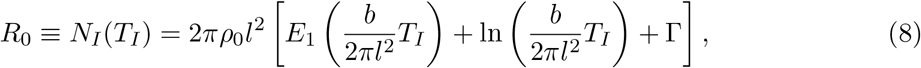

referred to as approximation 3. A comparison of the three approximations presented here, (4), (6), and (8), is shown in Figure 4. Note that for all the above approximations, reductions in susceptibles were only taken into account due to secondary infections caused from the singular primary infection, and not due to any further infections through the epidemic process, thus making all the approximations an over-estimate of *R*_0_ - we discuss this further in Section 3.3. In the parameter ranges considered, there is little difference between approximation 1 and approximation 2. Both overestimate the value of *R*_0_ in comparison to approximation 3, (8), particularly at a short-range infectious length scale, i.e., for *l <* 50 for *b* = 10. For this reason, for the remainder of this manuscript we consider (8) only as an analytic approximation to *R*_0_ and further investigate its suitability to describe the ABM in Section 3.3.

**Figure 4:**
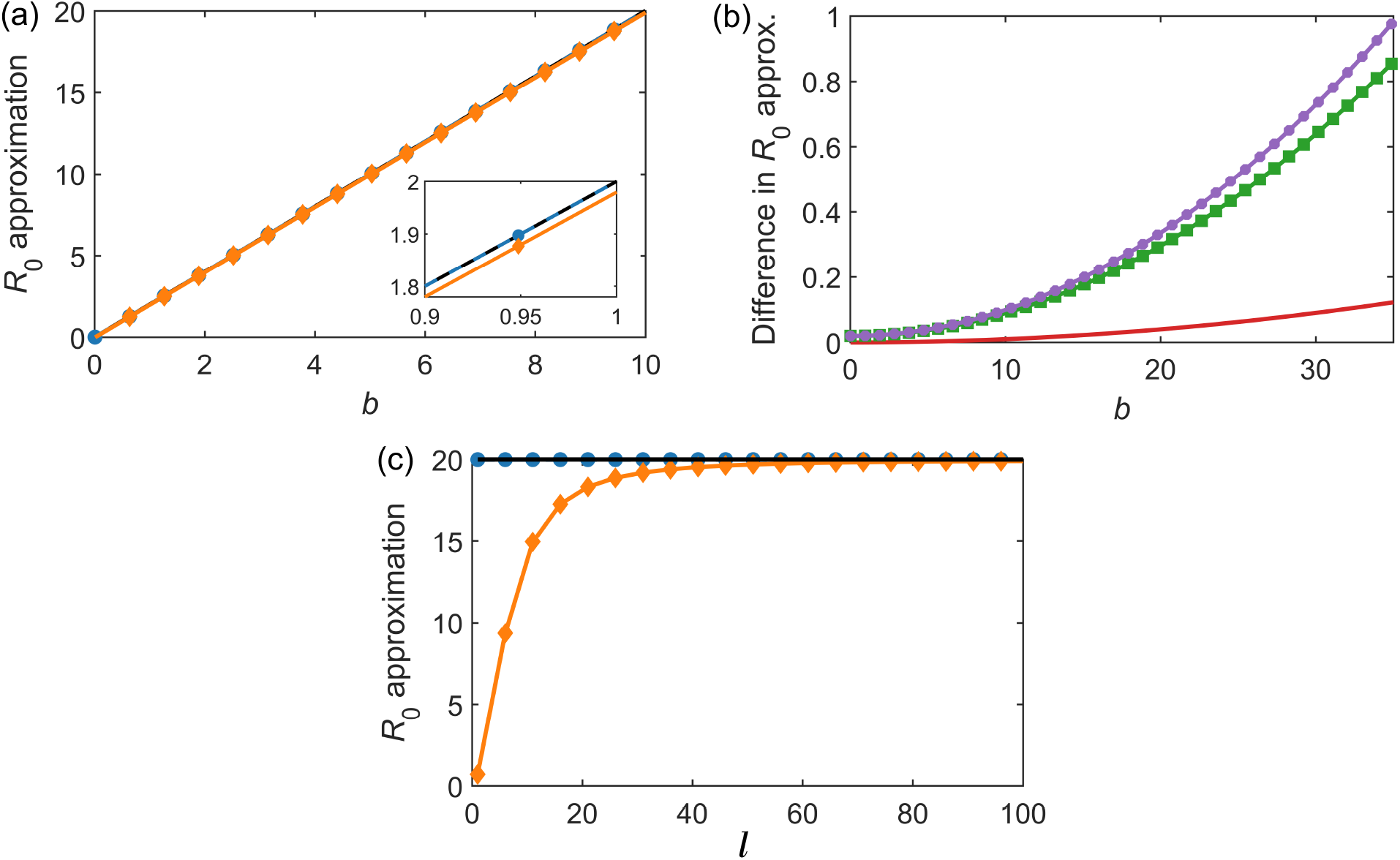
A comparison of the three analytic approximations to *R*_0_ presented in Section 2.5. (a) Approximation 1, (4), in black, approximation 2, (6), in blue dashed with circle markers and approximation 3, (8), in orange with diamond markers, for increasing values of infectious parameter *b*. The inset shows a smaller range of *b* to highlight the difference between the overlapping lines. (b) The differences in the *R*_0_ estimates between the three approximations, with the red solid line showing the difference between approximation 1 and 2, (4)-(6), the purple circle line showing the difference between approximation 1 and approximation 3, (4)-(8), and the green square line showing the difference between approximation 2 and approximation 3, (6)-(8). (c) Approximation 1, (4), in black, approximation 2, (6), in blue dashed with circle markers and approximation 3, (8), in orange with diamond markers, for increasing values of infectious length-scale parameter *l*.

## 3. Results

In this section we investigate the three methods of estimating the basic reproduction number from a typical agent-based model (ABM), described in Section 2. These methods include contact tracing (Section 3.1 using the methods described in Section 2.2.1), fitting to the standard SIR equations (Section 3.2 using the methods described in Section 2.4), and an analytical approximation (Section 3.3, using the methods described in Section 2.5). We then compare these methods by applying them to a simulation of the ash dieback pathogen in the UK (Section 3.4).

### 3.1. Contact tracing through the ABM

In the ABM described in Section 2.2, we assume infections can occur due to pressure from multiple sources through the infectious kernels surrounding each infected tree, as shown in Figure 2. This results in infection dynamics that are not one-to-one. We thus use the method for proxy-contact tracing described in Section 2.2.1, resulting in a time-step estimate for the (instantaneous) effective reproduction number, *R*(*t*). An example estimate of *R*(*t*), the corresponding instantaneous basic reproduction number calculated as *R*_0_(*t*) = *R*(*t*)*S*(0)*/S*(*t*), and a point estimate of *R*_0_ (calculated as the mean of *R*_0_(*t*)) for a single illustrative simulation are shown in Figure 5.

**Figure 5:**
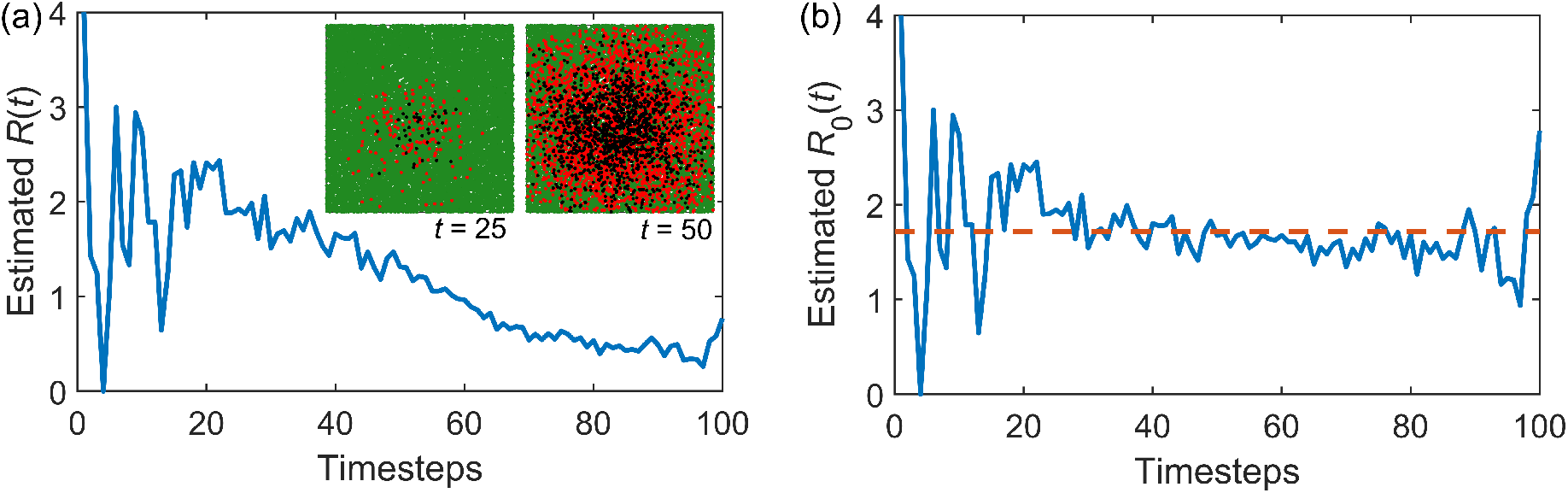
Proxy contact tracing through the ABM (with parameters *b* = 10, *l* = 100, *I*_0_ = 5, *L* = 1000, *ρ* = 0.02 and *T*_*I*_ = 10) to estimate (a) the instantaneous effective reproduction number, *R*(*t*), with snapshots of the disease spread with susceptible (green), infected (red) and removed (black) trees at *t* = 25 and *t* = 50 in the inset, in a bounded box of *L* = 1000 and (b) the instantaneous basic reproduction number *R*_0_(*t*) (blue solid line) and basic reproduction number *R*_0_ (orange dashed line, calculated as the mean of *R*_0_(*t*)).

We can consider how varying the two ABM infection parameters, the infectivity *b* and the length scale *l*, will impact the reproduction number, summarised in Figure 6 and Figure 7. The mean infected population time series (averaged over 500 ABM simulations for each parameter set) show the epidemic dynamics for increasing values of *b* and *l*, shown in Figure 6(a) and Figure 7(a), respectively. The mean instantaneous effective reproduction number *R*(*t*) for increasing *b* and *l* are shown in Figure 6(b) and Figure 7(b), respectively. As the ‘strength’ of the epidemic increases (through an increasing *b*, or to a lesser extent, an increasing *l*) the early values of *R*(*t*) increase. This results in a faster decreasing pool of susceptibles, and thus a more significant decrease in *R*(*t*) with time. The instantaneous basic reproduction number, *R*_0_(*t*), takes into account this decrease in susceptibles, and is shown for increasing *b* and *l* in Figure 6(c) and Figure 7(c), respectively, showing the expected increase in *R*_0_(*t*) with both increasing *b* and *l*. A single value of the basic reproduction number, *R*_0_, can be calculated as the mean of the instantaneous basic reproduction number for each simulation, shown in the box plots in Figure 6(d) and Figure 7(c) for increasing *b* and *l*, respectively. For increasing infectivity *b, R*_0_ increases linearly. For increasing length-scale *l, R*_0_ increases but saturates due to the presence of the bounding box (here at *L* = 1000), artificially decreasing the number of infections.

**Figure 6:**
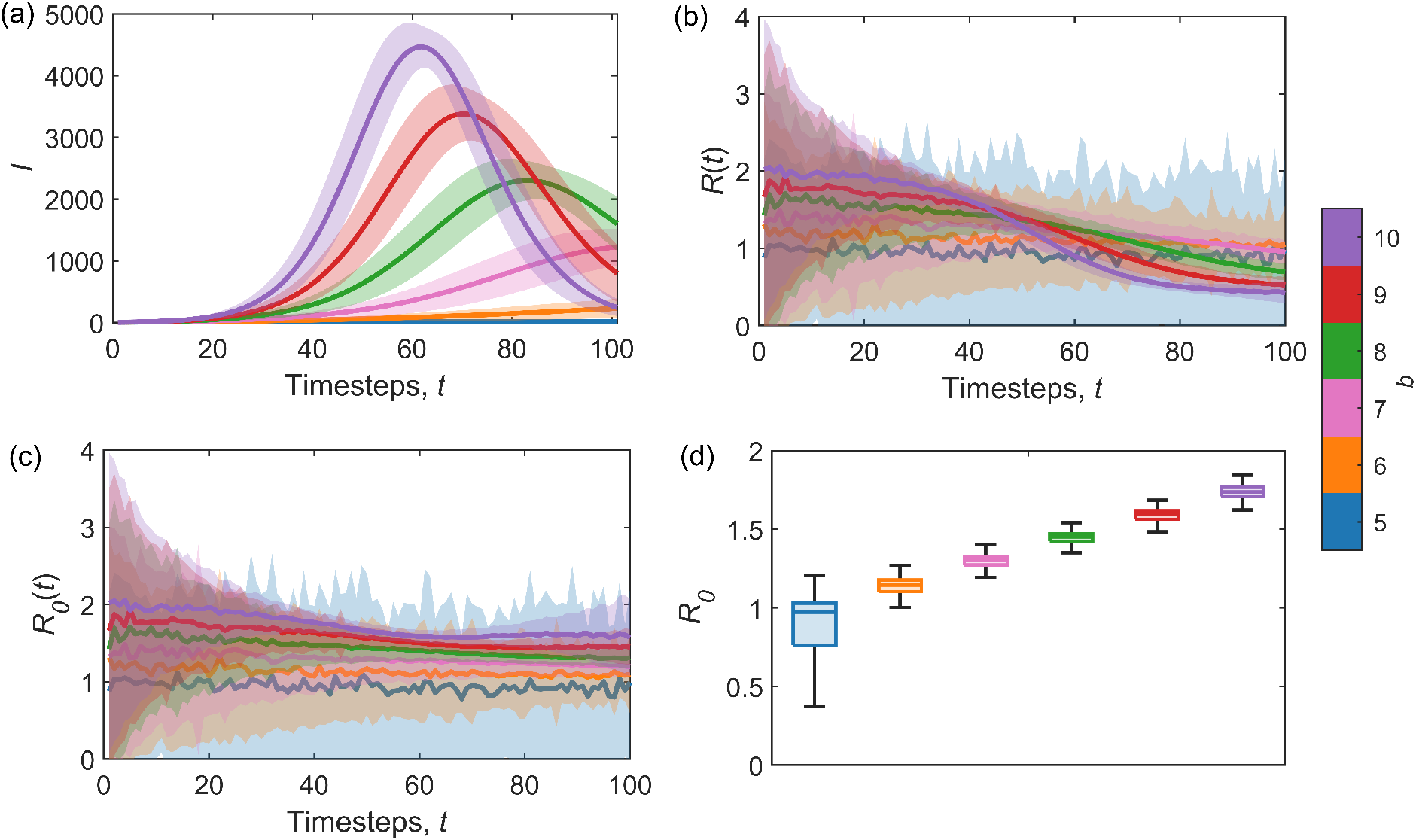
Proxy contact tracing through the ABM (with parameters: *l* = 100, *I*_0_ = 5, *L* = 1000, *ρ* = 0.02 and *T*_*I*_ = 10 at increasing values of infectivity *b*) to estimate *R*_0_ for varying *b*. The mean (from 500 simulations) (a) infected population with time, along with (b) the instantaneous effective reproduction number *R*(*t*) and (c) the instantaneous basic reproduction number *R*_0_(*t*). Error bars show the standard deviation. (d) Box plots of the estimated *R*_0_ values from each of the 500 runs (calculated as the mean of each of the 500 corresponding *R*_0_(*t*) time series).

**Figure 7:**
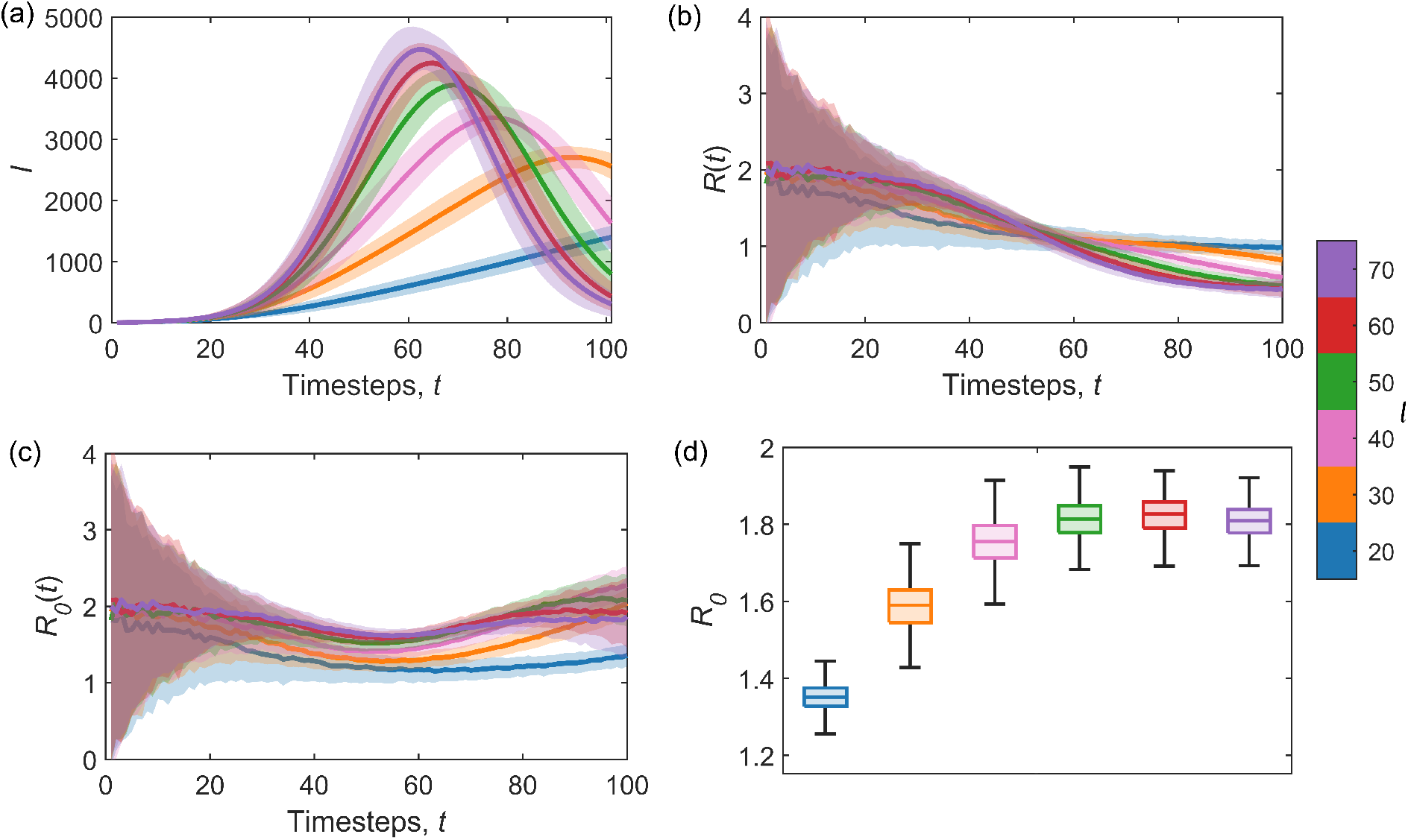
Proxy contact tracing through the ABM (with parameters: *b* = 10, *I*_0_ = 5, *L* = 1000, *ρ* = 0.02 and *T*_*I*_ = 10 at increasing values of infectious length scale *l*) to estimate *R*_0_ for varying *b*. The mean (from 500 simulations) (a) infected population with time, along with (b) the instantaneous effective reproduction number *R*(*t*) and (c) the instantaneous basic reproduction number *R*_0_(*t*). Error bars show the standard deviation. (d) Box plots of the estimated *R*_0_ values from each of the 500 runs (calculated as the mean of each of the 500 corresponding *R*_0_(*t*) time series).

### 3.2. Fitting the stochastic SIR equations

It may be necessary to avoid the computational expense of tracking infections through the ABM. In this case, we can compare the simulations to the SIR model [11] described with introduced stochasticity by (2) and (3). In this formulation, *R*_0_ is defined as *βN/γ*. Here, we employ the inference scheme detailed in Section 2.4 and Appendix A to estimate the SIR model parameter *β*. Since *γ* is known from the ABM input parameters (*γ* = 1*/T*_*I*_), we can then calculate *R*_0_. Example fittings of the stochastic SIR model for three characteristic spread dynamics with increasing dispersal length scale *l* are shown in Figure 8. At shorter length scales, we can see the slower increase in the infectious population, due to the spatial structure and a loss of the homogeneous mixing assumption when the interactions are short-range, particularly visible at the start of the epidemic due to a restricted number of susceptibles falling within the infection kernel. An advantage of this technique is the ability to obtain a posterior distribution of plausible *R*_0_ values (through the posterior distribution for *β* and our estimate of *γ* = 1*/T*_*I*_ from the ABM), shown in Figure 8 for each of the example infection spread dynamics.

**Figure 8:**
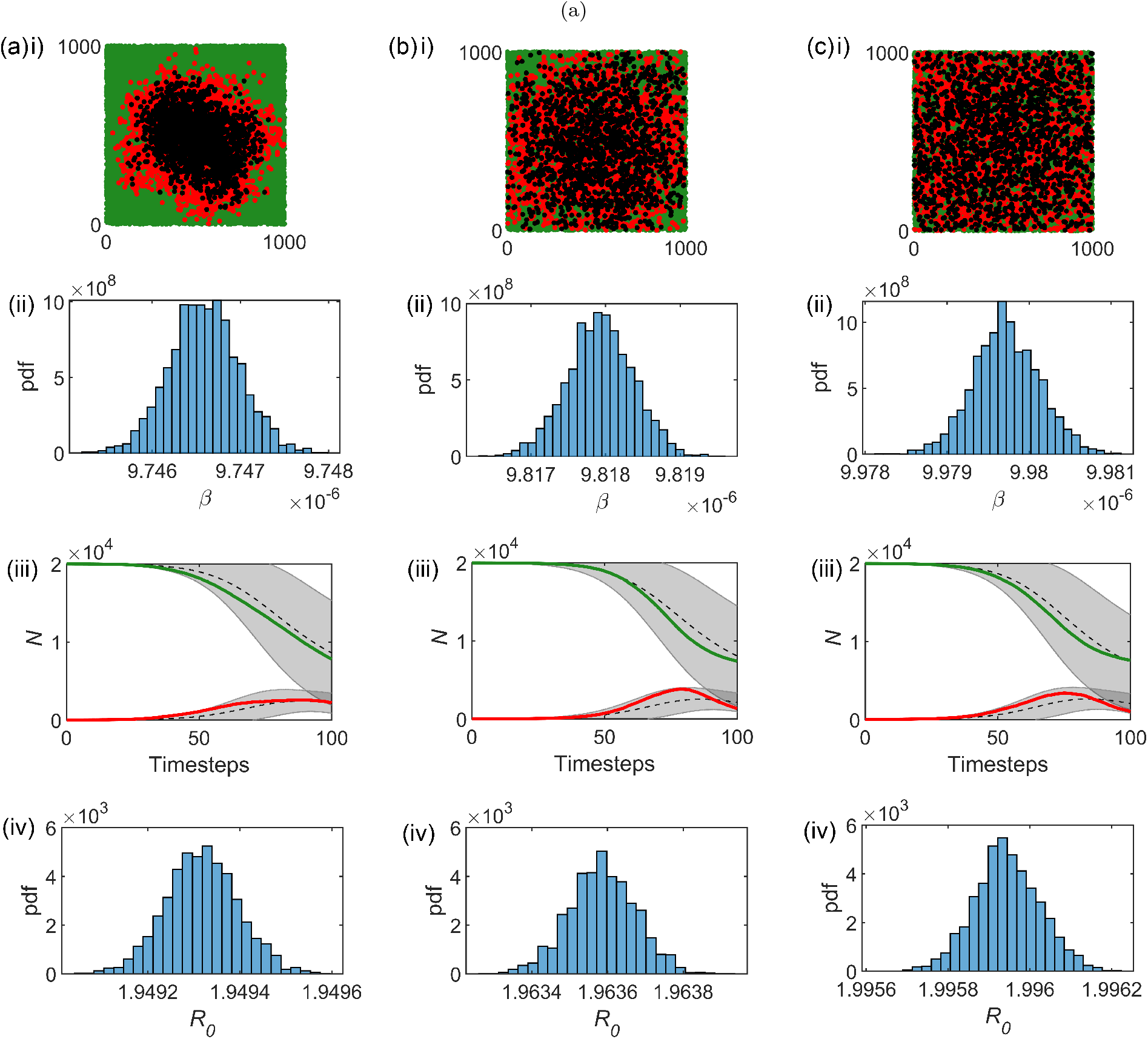
Example epidemic regimes (with *ρ* = 0.02, *T*_*I*_ = 10 and *L* = 1000) taking place over similar time scales with an increasing radius of infection *l*: (a) *b* = 10 and *l* = 40, (b) *b* = 10 and *l* = 100, and (c) *b* = 15 and *l* = 400, showing i) a snapshot at 20% mortality with susceptible (green), infectious (red) and removed (black) trees, ii) posterior distributions of the inferred SIR parameter *β*, (iii) the number of infectious (I, red) and susceptible (S, green) trees from the ABM, with the corresponding stochastic SIR fitting using the median inferred *β*, and *γ* = 1*/T*_*I*_ = 0.1, shown as the mean of 500 runs in black dashed with standard deviation error bars in grey) and iv) posterior distribution of estimated *R*_0_(*t*), using all posterior estimates of *β*, and *γ* = 1*/T*_*I*_ = 0.1.

### 3.3. An analytical approximation

An idealised, spatially explicit analytical approximation for *R*_0_ is derived in Section 2.5. In this section, we compare the analytic predictions of *R*_0_ resulting from (8) to the numerical results from proxy contact tracing within the ABM (as described in Section 2.2.1 and Section 3.1).

The cumulative number of expected secondary infections due to a primary infection over an infectious period of *T*_*I*_ = 100, *N*_*I*_(*t*), is shown in Figure 9(a) for a fixed dispersal parameter *l* = 100 and increasing *b*. The linear relation between time and *N*_*I*_(*t*) was predicted by (8) in Section 2.5). For lower values of *b*, the analytic approximation captures the estimation from the proxy contact tracing. For *b* = 8, the analytic estimate from (8) overestimates *N*_*I*_(*t*) from around *t* = 50, as it only takes into account a reduction in susceptibles from infections caused by the primary infectious individual, and not other infections involved in the epidemic process. The contact-traced estimate shows the actual saturation of *N*_*I*_(*t*) that occurs as the number of available susceptibles limits the infection spread. Similarly, Figure 9(b) shows *N*_*I*_(*t*) for fixed *b* = 5 and increasing dispersal parameter *l*. In this case, the overestimation from the analytic *N*_*I*_(*t*) occurs at shorter length scales (e.g., for *l* = 10 in this parameter regime), where the pool of susceptibles has been reduced in a small area around the primary infection, limiting the epidemic process, as shown by contact tracing. A larger dispersal kernel encompasses a larger local neighbourhood of infectivity around the primary susceptible, resulting in less sensitivity to reductions in susceptibles through subsequent infections.

**Figure 9:**
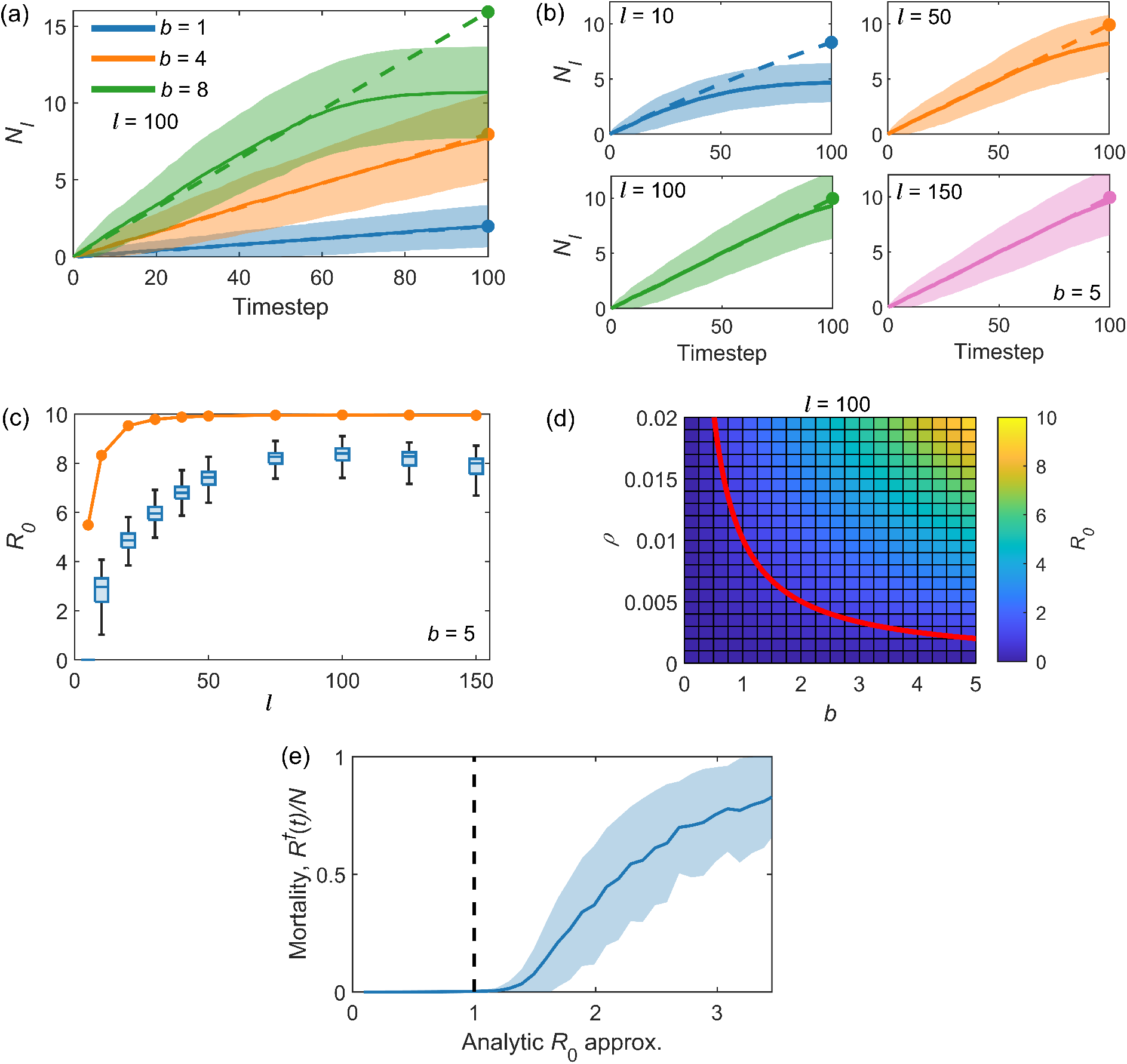
Comparison between the analytical expression for *N*_*I*_ (*t*) (the cumulative number of secondary infections resulting from the primary infection), and hence *R*_0_ = *N*_*I*_ (*T*_*I*_) from (8), and the value of *R*_0_ estimated through proxy contact tracing through the ABM (as in Section 3.1). In all cases *L* = 1000, *ρ* = 0.02, *I*_0_ = 1 and *T*_*I*_ = 100. (a) *N*_*I*_ for increasing infectivity *b* (with fixed infectious length scale *l* = 100). The analytic estimate of *N*_*I*_ (*t*) (from (8)) is shown as a dashed line, with a circle at the end of the infectious period indicating *R*_0_. The solid lines show the median contact traced estimates of *R*_0_ (over 500 runs of the ABM) with standard deviation error bars. (b) *N*_*I*_ for increasing infectious length scale *l* (with fixed infectivity *b* = 5). The analytic estimate of *N*_*I*_ is shown as a dashed line, with a circle at the end of the infectious period indicating *R*_0_. The solid lines show the median contact traced estimates of *R*_0_ (over 500 runs of the ABM) with standard deviation error bars. (c) The analytic estimates of *R*_0_ for increasing length scale *l* (and fixed infectivity *b* = 5) is shown as the solid orange line with circle markers. Box plots show the corresponding estimates of the contact traced *R*_0_ values from 500 runs of the ABM. (d) The analytic *R*_0_ phase plane predicted by (8) for increasing density *ρ* and infectivity *b*. The threshold, given by *R*_0_ = 1, is plotted in red and illustrates the separation between confinement and epidemic. (e) The relationship between the total tree mortality (removal prevalence) over 500 runs of the ABM and the *R*_0_ value predicted by (8), demonstrating the threshold-like behaviour at *R*_0_ = 1.

The approximation of the basic reproduction number *R*_0_ is given by the cumulative number of secondary infections resulting from the primary infection over its infectious period, i.e., *R*_0_ = *N*_*I*_(*T*_*I*_). In general, (8) overestimates *R*_0_ due to the neglect of any subsequent infections caused by the secondary infections. This is exacerbated when the infectivity is high, or the length scale of dispersal is low, as seen in the plateau of infections in Figure 9(a) and (b). Thus, (8) describes constant transition rates accurately but deviates from model simulations when subsequent infections cause a significant reduction in local susceptible density. The values of *R*_0_ estimated through both the analytic and numeric contact tracing approximations for an arbitrary fixed infectivity of *b* = 5 and an increasing dispersal parameter *l* are shown in Figure 9(c). The analytic approximation captures the general trend of increasing *R*_0_, followed by saturation due to a limited susceptible population in the infectious area, as in Figure 7(d). An increase in the size of the ABM area (the box defined by *L*) would shift this saturation to larger length scales. A useful property of the basic reproduction number is its quantification of the transmission threshold, defined as *R*_0_ = 1, predicting the separation of states between confinement and epidemic. In Figure 9(d), the threshold predicted by (8) is shown with a two-dimensional phase plot of all estimated *R*_0_ over tree density and infection parameter *b*, allowing a categorisation of disease confinement or epidemic from the model parameters.

We can also assess how the total tree mortality relates to the threshold *R*_0_ > 1 predicted by (8). The total proportion of host trees in the Removed (*R*^*†*^) compartment (removal prevalence) for increasing predicted analytical *R*_0_ from (8) is shown for an illustrative epidemic regime (*L* = 500, *ρ* = 0.01, *T*_*I*_ = 100, *r* = 100, 0 *< b <* 4) in Figure 9(e). When *b* and *ρ* result in an analytical prediction of *R*_0_ less than one, correspondingly, tree mortality is low. When the infectivity is increased such that analytical *R*_0_ > 1, tree mortality rises considerably. The approximation described by (8) therefore demonstrates the threshold-like behaviour defined by *R*_0_ = 1. It is worth noting that despite being above the threshold, the numerical simulations from the ABM can still fail to produce an epidemic, due to the influence of early stochastic forces increasing the probability of epidemic extinction [25, 26], a disadvantage of the general concept of *R*_0_ [27]. The threshold-like behaviour witnessed in Figure 9(e) demonstrates that (8) provides a simple predictive framework for the ABM considered here.

### 3.4. Application to ash dieback

In this section we employ the idealised ABM, as described in Section 2.2, to describe the spread of ash dieback (*Hymenoscyphus fraxineus*), a highly destructive fungal tree disease, and use each of the above methods to estimate the basic reproduction number *R*_0_.

The ABM is initialised with a density of *ρ* = 0.0444 trees/m^2^, given the estimate of 444 ash trees per hectare (10,000 m^2^) within small woodlands in the UK [28]. We set the dispersal parameter *l* in (1) to be 138 m, based upon estimates of the local dispersal kernels for the ash dieback pathogen from spore-trapping data [10]. The pathogen can remain active for around 4 years after infection [29] and so the infectious period is chosen to be *T*_*I*_ = 4 years. This leaves the unconstrained infectivity parameter, *b*, and the time-step over which new infections are assessed, *dt*. Here we arbitrarily take *dt* = 1 week, and consider a range of *b* (with units *m*^2^week^*−*1^) to focus on regimes resulting in 0 *< R*_0_ < 7.

Assuming there will be some variability in the estimate of *ρ* in different areas, we show the phase plane of the analytic estimation of *R*_0_ for a range of densities and infectivity parameters, along with the threshold value of *R*_0_ = 1, in Figure 10(a). This illustrates the values of *ρ* and *b* that would result in the epidemic regime. The estimates of *R*_0_ for 500 simulations with fixed *ρ* = 0.0444 trees/m^2^ and *l* = 138 m through each of the methods described above is shown in Figure 10(b). The overestimate of *R*_0_ through analytical means is clear, as discussed in Section 3.3, with similar estimates of *R*_0_ resulting through both contact tracing and inference of the SIR parameters. Given an estimate of the infectious parameter *b* (discussed further in Section 4), this would allow the prediction of *R*_0_ through several comparative methods without requiring physical contact tracing of the infection.

**Figure 10:**
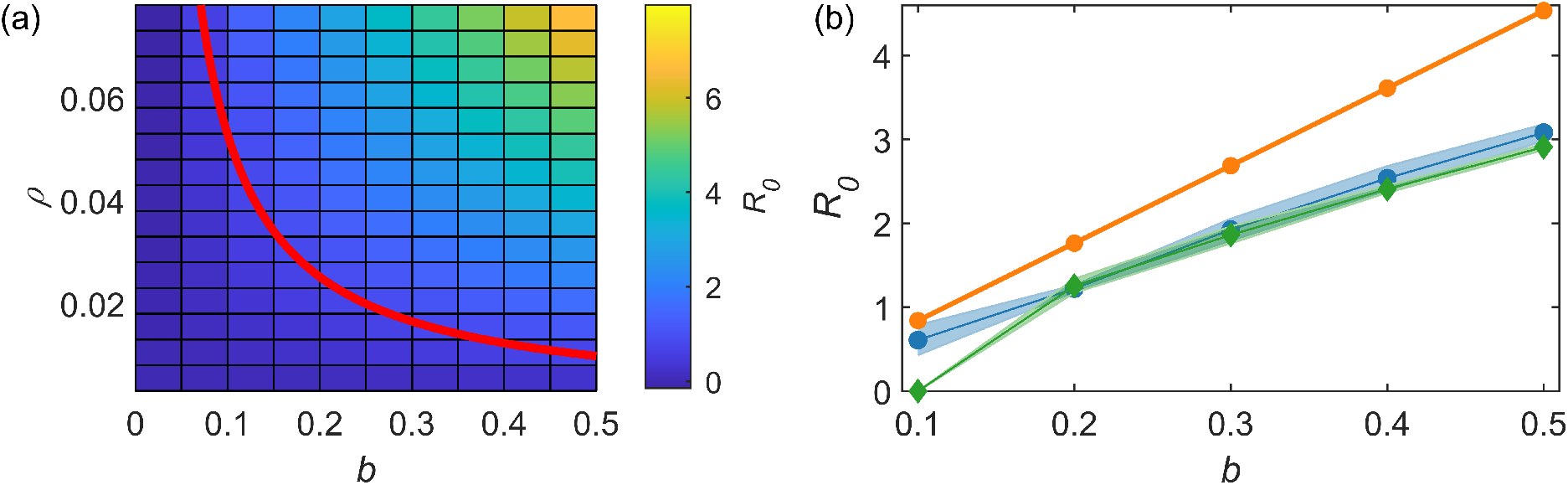
(a) The analytic *R*_0_ phase plane predicted by (8) for increasing density *ρ* (including the estimated tree density relevant for ash dieback, *ρ* = 0.0444) and infectivity *b*. The threshold, given by *R*_0_ = 1, is plotted in red and illustrates the separation between confinement and epidemic. (b) The estimates of *R*_0_ through the analytic approximation (8) (solid line with orange circles), contact tracing where *R*_0_ is the mean of *R*_0_(*t*) (blue circles), and inference of the SIR parameters (green diamonds). Markers indicate the mean of 500 simulations, with standard deviation error bars.

## 4. Discussion

Stochastic ABMs are a popular choice for describing the spread of tree diseases and pests through woodland areas, due to their flexibility and ability to describe large-scale collective behaviours from the programming of individual behaviours. Here we use a typical ABM for the spread of an arbitrary disease or pest through a synthetic forest, and explore different methods of calculating the basic reproduction number *R*_0_, a key parameter for disease characterisation and forecasting.

We have expanded upon similar compartmental ABMs of tree disease [7], introducing a flexible spatial component to the infection spread through the inclusion of an infectious kernel, as used in [9] to describe the spread of Asiatic citrus canker disease. The infectious dispersal kernel characterises the spatial spread of the disease and thus has a large impact on the epidemic outcome [30]. In this case we use an ABM with a Gaussian probability of infection due to its simplicity and prevalence in similar models [19, 16, 30, 31], but this is entirely flexible and the most appropriate choice will depend on the nature of the disease or pest considered. For example, for the ash dieback epidemic, both Gaussian and power law kernels have been estimated for the dispersal at different scales [10]. Future work should explore the impact of different kernels on the ABM behaviour and the corresponding estimations of *R*_0_.

The choice of infectious kernel is linked to dispersal [19, 32] describing the probability of pathogen presence over distance. Dispersal kernels can be estimated through either mechanistic models [33, 34], or through fitting statistical functions to dispersal data [32, 10], but can be challenging to ascertain. A further complication is the mapping of a dispersal kernel to the corresponding infectious kernel, i.e., how does probability of pathogen presence correspond to the probability of successful infection? In the ABM kernel this is captured in the infectivity parameter *B* = *b/*(2*πl*^2^), and we consider a range of arbitrarily chosen values to illustrate the methodology. For the ash dieback simulation, we take the length scale from the dispersal kernel estimated from ecological data describing the density of pathogen spores at varying distance from infected ash trees [10]. Since *B* (and thus *b*) is not known, we again consider a range of parameter values leading to plausible *R*_0_ estimations.

Inference for *B* under the ABM may be possible by leveraging recently proposed approximate Bayesian computation techniques (ABC, see e.g. [35]) although such approaches typically require several millions of model simulations. The practical applicability of ABC for our proposed spatio-temporal model remains the subject of ongoing work. It may also be possible to constrain *B* through estimations of *R*_0_ (through empirical means or from data of infections) in previous outbreaks of the disease or pest under similar conditions. Climate change is driving the geographical expanse of many ecological epidemics and invasions, and so ecological data may be available from multiple locations.

The most straightforward way of calculating *R*_0_ within the ABM is through the contact tracing of infections. This can be done directly, by storing the number of secondary infections resulting from a singular tree, but requires a model in which trees are considered individually in the infection time-step, thus requiring more computational expense. In the case of a vectorised ABM, where individual secondary infections are not tree specific, we can calculate an estimation of *R*(*t*) through the number of average new infections, and sharing ‘the blame’ for these between the currently infected population. Although noisy, particularly at the beginning of an epidemic regime (if the length scale of infectivity is low and initial infected trees are clustered, resulting in a limited number of susceptible trees in the local area) and end of epidemic simulation (as the number of susceptible trees available approaches zero), the median of the *R*_0_(*t*) series provides an estimation consistent with the other methods considered here. Contact-tracing methods have the advantage of being simple to implement computationally, even with increasing model complexity.

The temporal output from the ABM can be compared to the classic SIR equations, allowing a straight-forward estimation of *R*_0_ through the model parameters *β* and *γ*. The estimation of the model parameters through the inferential scheme results in a distribution of plausible *R*_0_ values, capturing the parameter uncertainty and useful for forecasting best and worst-case scenarios. This method relies on the assumption that the standard SIR equations will effectively capture the time-series resulting from the ABM. Unless the length scale of dispersal is very large, we will not fulfil the homogeneous mixing assumption of the SIR model, however we still find the parameter estimates to be descriptive enough to provide a measure of *R*_0_ consistent with other methods. Future work could explore the relationship between the dispersal kernel parameters and the SIR model parameters.

We also derive an analytical expression for approximating *R*_0_ which predicts the epidemic threshold in line with the contact-traced reproduction number computed through the ABM simulations, with the caveat that it overestimates *R*_0_, particularly when the epidemic severity is high. The overestimation of *R*_0_ can be compared to well-known results [26, 36] showing that the first-generation basic reproduction number for farms infected with foot-and-mouth overestimates the growth rate of infection. This estimate has the advantage of calculation through the ABM model parameters only, however comparable analytical solutions are challenging to determine for more elaborate life-cycles, dynamics and aggregated host distributions.

Considering an illustrative simulation of ash dieback in the UK using the ABM allows a comparison of the different estimation methods of *R*_0_ in context. If the infectivity parameter *b* were to be estimated (through estimation of *B* and *l*), as discussed above, the ABM model parameters could be used to efficiently calculate an upper-bound for a plausible estimation of *R*_0_. Numerical methods such as contact tracing and inference of the SIR model parameters provide a more accurate estimation of *R*_0_, albeit at more computational expense. Future work could expand the ABM to larger areas, considering varying densities of woodland across different geographical areas to generate landscape level predictive *R*_0_ maps.

This work outlines a framework for estimating epidemic severity through the parameter *R*_0_, using a flexible ABM for tree disease spread. Given knowledge of the dispersal kernel for a particular pathogen, the ABM can be used to estimate *R*_0_ through several methods, and thus provides an extensible tool which can be further developed for ecological epidemic forecasting.

## Acknowledgements

This research was supported by: EPSRC New Horizons Grant EP/V048511/1 (AB, AG, NGP, and LW), NERC Knowledge Exchange Fellows Grant NE/X000478/1 (LW) and DEFRA UK (JH, JS, and NGP).

## Appendix A Bayesian inference details

For inference on the stochastic SIR model, (2) and (3), we apply the linear noise approximation (LNA) to obtain a tractable approximation to the stochastic differential equation. Formal details of the LNA can be found in [37, 38, 39], with an outline derivation below.

Consider a partition of the system *X*_*t*_ as

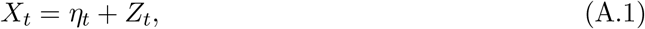

where *{η*_*t*_, *t ≥* 0*}* is a deterministic process satisfying the ordinary differential equation (ODE)

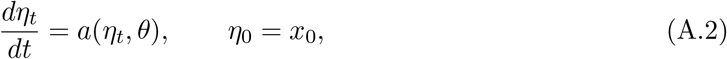

and *{Z*_*t*_, *t ≥* 0*}* is a residual stochastic process. The residual process *Z*_*t*_ satisfies

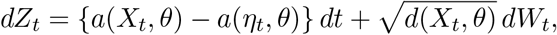

which will typically be intractable. The assumption that ||*X*_*t*_ *− η*_*t*_|| is “small” motivates a Taylor series expansion of *a*(*x*_*t*_, *θ*) and *d*(*x*_*t*_, *θ*) about *η*_*t*_, with retention of the first two terms in the expansion of *a* and the first term in the expansion of *b*. This gives an approximate residual process 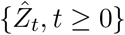 satisfying

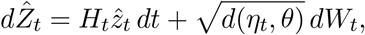

where *H*_*t*_ is the Jacobian matrix with (*i,j*)th element

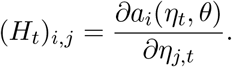

For the SIR model in (2) and (3) we therefore have

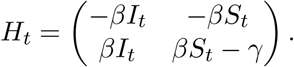

Given an initial condition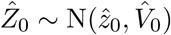, it can be shown that 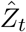 is a Gaussian random variable [40]. Consequently, the partition in (A.1) with *Z*_*t*_ replaced by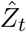, and the initial conditions *η*_0_ = *x*_0_ and 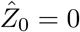 give

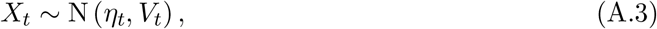

where *η*_*t*_ satisfies (A.2) and *V*_*t*_ satisfies

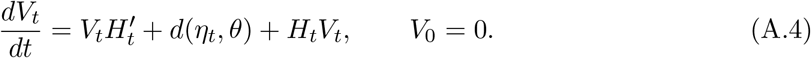

Further details on the derivation of (A.4) are given in [17]. Hence, the linear noise approximation is characterised by the Gaussian distribution in (A.3), with mean and variance found by solving the ODE system given by (A.2) and (A.4), which can be solved numerically.

Given the observational process *X*_*t*_, and parameter vector *θ*, the likelihood is then

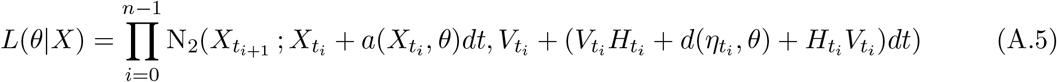

where N_2_(*·, m, v*) denotes the multivariate Gaussian density with mean vector *m* and variance matrix *v*. In this case we fix *γ* according to the removal rate in the ABM and so *θ* = (*β*) only. We set prior specifications of *β ∼* logN(0, 1). Since *β >* 0 we work on a unrestricted parameter space by letting *λ* = log *β*. The posterior is given by

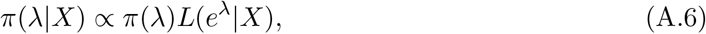

where 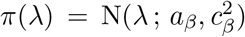 We can then use an MCMC scheme (Algorithm 1) to generate draws of *λ*|*X* and exponentiate to give draws of *θ*|*X*. This results in a posterior distribution of plausible values of *β*. In cases where *γ* is not known, it is straightforward to expand the parameter search to target *θ* = (*β, γ*)′.

### Algorithm 1

Random walk Metropolis algorithm

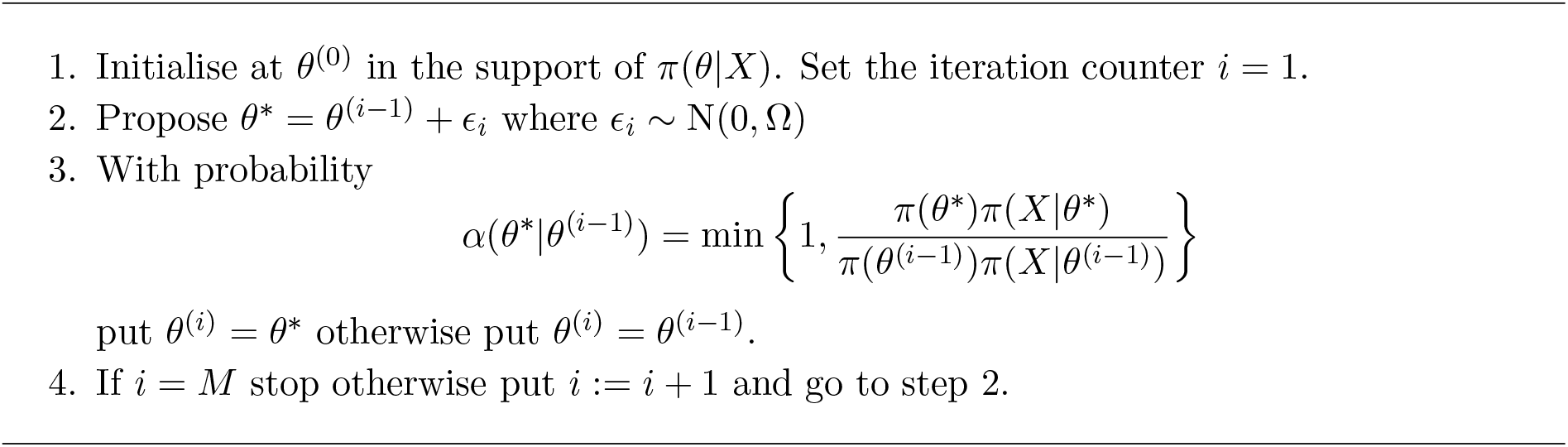

## Appendix B. Analytic *R*_0_ derivation

In this section, an idealised, spatially explicit expression of *R*_0_(*t*) is derived for the ABM. Firstly, consider a single infectious individual with infectious period *T*_*I*_, surrounded by susceptibles at varying distance *r* at a density *ρ*. We can estimate the number of new infections caused by the infected individual to be equal to the number of susceptibles at distance *r, S*(*t*) = 2*πrρ*, multiplied by the probability of infection at *r*, in this case described by (1). The expected number of new secondary infections from the initial infected individual at time step *t* (for 1 *< t < T*_*I*_) for all possible values of *r* is therefore

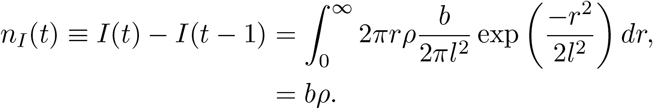

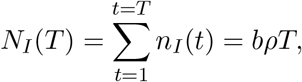

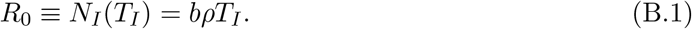

However, this assumes that the density of the susceptibles remains fixed at each time step as the infection process progresses.

If host growth is neglected (likely appropriate for many tree based scenarios), fewer trees will be available to infect at time-step *t* + 1. The susceptible tree density can therefore be seen as a monotonically decreasing function of time, *ρ*(*t*), leading to

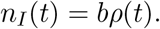

In a large but finite domain of size *L*, tree density approximately follows

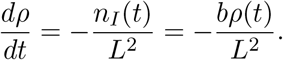

Solving the above, with the initial condition *ρ*_0_ at *t* = 0, leads to

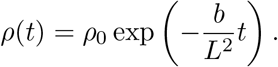

The cumulative sum of new secondary infections, during a set time period 0 ≤ *t* ≤ *T*, is then given by

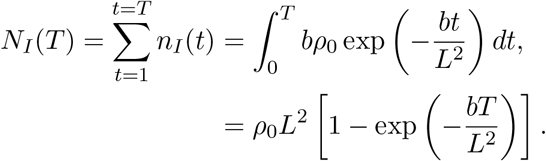

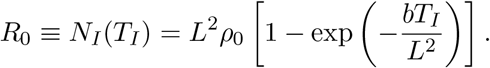

This can be considered as a second approximation to *R*_0_ where some reduction in the number of susceptibles has been taken into account. Note that this is an idealised scenario in which density reductions due to secondary and tertiary infections are neglected.

However, the uniform density reductions in the above assume that secondary infections are equally likely at all spatial locations about the primarily infected tree, which is not the case for an infectivity kernel as in (1). On average, neglecting this spatial variation within the changing density results in an overestimation of the number of secondary infections induced by the tails of the dispersal kernel, thus giving rise to a greater *R*_0_ value. Taking this into account, we can instead consider a spatial variant density of the form

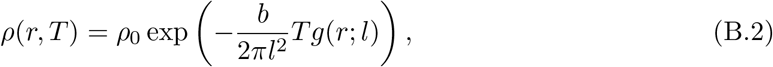

where *g*(*r*; *l*) is a Gaussian kernel as in (1). As above, the total number of new secondary infections from the primary infection during a set time period, 0 ≤ *t* ≤ *T*, at a certain distance *r, N*_*I*_(*r, T*), can be calculated from the overall change in density

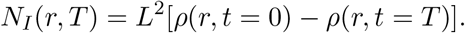

For all possible values of *r* this becomes

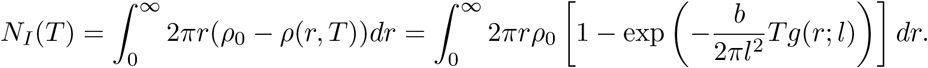

Here the finite lattice square of area *L*^2^ has been replaced with integration in polar coordinates over *dr*. Although the above is difficult to solve directly, it can be integrated by performing a series expansion on the exponential term and integrating on a term-by-term basis, as follows:

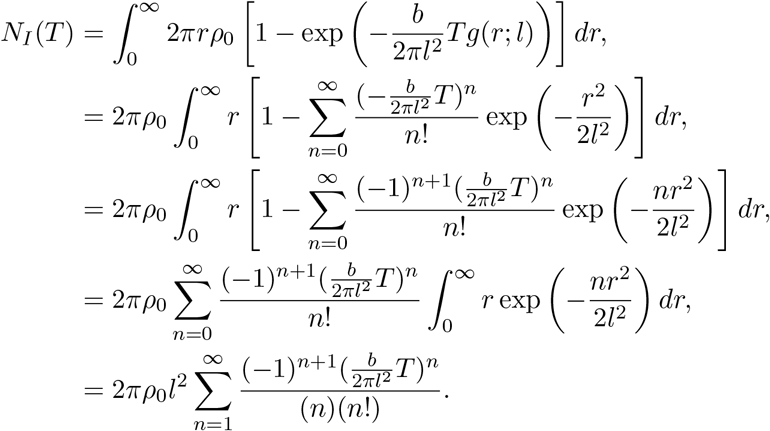

If *b/*2*πl*^2^ and *T* are small, the first order term in the above equation is sufficient to approximate *N*_*I*_(*t*) as a linear function of *T*. Finally, the above can be summed to give

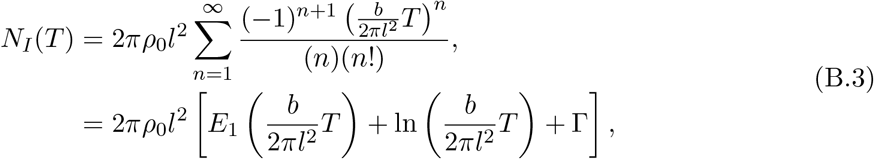

where the function *E*_1_(*x*) is the mathematically well studied exponential function 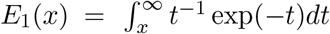 and Γ is the Euler-Mascheroni constant *≈* 0.57721. The estimated value of the basic reproduction number is therefore

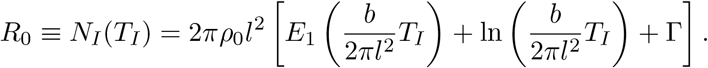

